# Appearance of synthetic vector-associated antibiotic resistance genes in next-generation sequences

**DOI:** 10.1101/392225

**Authors:** George Taiaroa, Gregory M. Cook, Deborah A Williamson

**Author notes:** Email addresses.

## Abstract

**Background:** Next-generation sequencing methods have broad application in addressing increasing antibiotic resistance, with identification of antibiotic resistance genes (ARGs) having direct clinical relevance.

**Objectives:** Here, we describe the appearance of synthetic vector-associated ARGs in major public next-generation sequence data sets and assemblies, including in environmental samples and high priority pathogenic microorganisms.

**Methods:** A search of selected databases – the National Centre for Biotechnology Information (NCBI) nucleotide collection, NCBI whole genome shotgun sequence contigs and literature-associated European Nucleotide Archive (ENA) datasets, was carried out using sequences characteristic of pUC-family synthetic vectors as a query in BLASTn. Identified hits were confirmed as being of synthetic origin, and further explored through alignment and comparison to primary read sets.

**Results:** Synthetic vectors are attributed to a range of organisms in each of the NCBI databases searched, including examples belonging to each Kingdom of life. These synthetic vectors are associated with various ARGs, primarily those encoding resistance to beta-lactam antibiotics and aminoglycosides. Synthetic vector associated ARGs are also observed in multiple environmental meta-transcriptome datasets, as shown through analysis of associated ENA primary reads, and are proposed to have led to incorrect statements being made in the literature on the abundance of ARGs.

**Conclusions:** Appearance of synthetic vector-associated ARGs can confound the study of antimicrobial resistance in varied settings, and may have clinical implications in the nearfuture.

## Introduction

Elevated and increasing rates of antimicrobial resistance pose a significant challenge, in both medical and agricultural settings^1^. The identification of characterised antibiotic resistance genes (ARGs) can allow for the assessment of resistance status, and inform clinical practice^2^. Next-generation sequence-based data (reads, assemblies) are increasingly being used for the identification of ARGs, and for some pathogens, replacing phenotypic methods^2,3^. These data also enable broader study of ARG abundance and dissemination between organisms and geographic locations^4,5^. Synthetic ARGs have previously been shown to contaminate common laboratory reagents^6–9^, having the potential to confound PCR-based ARG identification in biological samples^8–10^. Here, we explore the appearance of synthetic ARGs in a range of next-generation sequence-based data, and comment on the impacts of this appearance in several previous studies.

## Methods

A detailed methods section can be found in the ‘Supplementary Materials’. In brief, an initial focus was placed on synthetic vectors of the pUC-family^11^, commonly used in molecular biology for the generation of laboratory reagents, and which we hypothesised to be a source of contamination in next-generation sequence-based data. Characteristic sequences located on the pUC-family of synthetic vectors provided the basis of a BLASTn search of two selected data sets: the NCBI nucleotide collection, consisting of GenBank, EMBL, DDBJ, PDB and RefSeq sequences but excluding expressed sequence tags (ESTs), (sequence-tagged sites (STSs), genome survey sequences (GSSs), whole-genome shotgun (WGS) sequences, transcriptome shotgun assembly (TSA) sequences, the Short Read Archive (SRA) and patent sequences, and a second search of the NCBI whole genome shotgun contigs (wgs) database. Characteristic sequences of the pUC-family initially included the pUC19 origin of replication region and beta-lactamase *bla_TEM_^11^*, and later the aminoglycoside-3’-phosphotransferase *aph(3).* Identified hits were confirmed as synthetic through analysis of associated genes and known synthetic vector structures. Samples that had associated metadata identifying the sequences as vector-containing, or otherwise transgenic, were excluded from analyses. Identified synthetic vectors were aligned, and associated ARGs additional to *blaTEM* identified. Literature examples of ARGs appearing in unexpected taxa were explored using primary read sets, and available assemblies where appropriate.

## Results and Discussion

### Synthetic vector-associated ARGs are attributed to organisms across the tree of life

Based on the above methods, synthetic vector-associated ARGs were identified in next-generation sequence-based data attributed to a broad range of organisms. Of those found in the NCBI wgs database, examples of all Kingdoms of life were represented. A summary is shown in Figure 1, as a phylogenetic tree including organisms of the Archaea, Eukarya and a selected branch of the Prokarya. Of those organisms identified, notable high priority pathogens^12^ include *Staphylococcus aureus, Clostridium difficile* and *Mycobacterium tuberculosis* (Table 1, Summary Table 1). Identified hits include biological samples prepared and sequenced between 2010-2018, employing a range of methods for next-generation sequence library preparation, sequenced on various platforms, and from diverse geographic locations (Supplementary Table 1).

**Figure 1.**
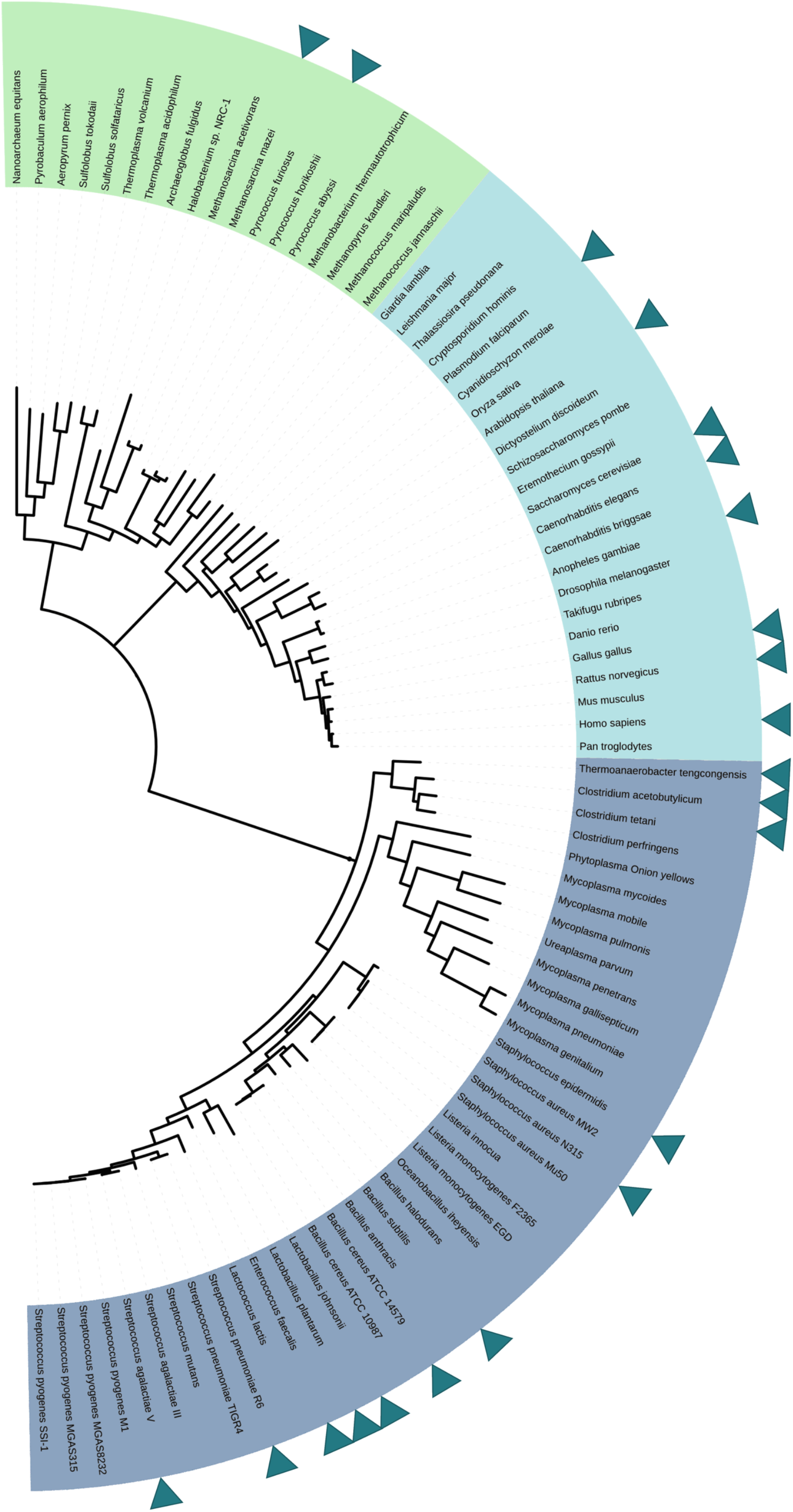
A simplified phylogeny showing Archaea (green), Eukarya (light blue) and the firmicute branch of the Prokarya (dark blue), with appearances of synthetic vector-associated ARGs carrying *blaTEM* highlighted by triangles beside species of the nearest taxa shown

**Table 1:** Examples of organisms entered in the NCBI wgs database with apparent synthetic vector associated ARGs

| Organism | Entry | Synthetic ARG/s | Country |
| --- | --- | --- | --- |
| *Bacillus cereus*<sup>a</sup> | 2016 | *blaTEM*, *tetC* | Switzerland |
| *Candida auris*<sup>b</sup> | 2015 | *blaTEM* | India |
| *Clostridium botulinum*<sup>c</sup> | 2017 | *aph(3')*, *blaTEM* | France |
| *Clostridium difficile*<sup>d</sup> | 2012 | *blaTEM* | United Kingdom |
| Corynebacterium ulcerans<sup>e</sup> | 2015 | *blaTEM, tetC* | Canada |
| Enterococcus sp.<sup>f</sup> | 2012 | *blaTEM* | USA |
| Fusarium oxysporum<sup>g</sup> | 2017 | *blaTEM* | Germany |
| Mycobacterium bovis<sup>h</sup> | 2018 | *blaTEM* | USA |
| Mycobacterium caprae<sup>i</sup> | 2018 | *blaTEM* | USA |
| Mycobacterium tuberculosis<sup>j</sup> | 2016 | *blaTEM, tetC* | United Kingdom |
| Peptoclostridium difficile<sup>k</sup> | 2016 | *blaTEM* | United Kingdom |
| Pseudomonas aeruginosa<sup>l</sup> | 2015 | *AAC(3), blaTEM* | Canada |
| Staphylococcus aureus<sup>m</sup> | 2015 | *blaTEM* | USA |
| Staphylococcus capitis<sup>n</sup> | 2018 | *blaTEM* | Canada |
| Staphylococcus epidermidis<sup>o</sup> | 2018 | *blaTEM* | Canada |

a. LQZZ01000237 b. CVRJ01000347 c. LIIS01000040
d. AHJJ01000181 e. JZUS01000009 f. AJQX01000216
g. FMJY01000040 h. MWXE01000433 i. MWXD01000140
j. FPPK01000048 k. FJOI01000053 l. LORB01000003
m. LALL01000105 n. PZCW01000079 o. PYYZ01000131

Similarly, in the NCBI nucleotide collection, synthetic-vector associated ARGs were found broadly, with examples of published genomes carrying these sequences including the wild tomato *Solanum pennellii* (HG975449), the red yeast *Xanthophyllomyces dendrorhous* (LN483254), and the tamar wallaby *Notamacropus eugenii* (CR388000). A subset of the hits in the NCBI nucleotide collection included samples sequenced using BAC cloning, which could be a possible non-contaminating source of synthetic sequences, and so are not considered further here.

### Synthetic vectors harbor a range of ARGs

As well as the targeted ARG blaTEM, numerous other synthetic vector-associated ARGs appear on identified contigs found in the NCBI wgs database (Supplementary Table 1). Frequently appearing ARGs include *aph(3’)* associated with resistance to aminoglycosides, and *tetC,* associated with resistance to tetracyclines. Less frequently appearing ARGs include *aac(3)* (aminoglycoside resistance), *blaCTXAM* (betalactam resistance), *ble* (bleomycin resistance), *catA1* (chloramphenicol resistance )and *ermB* (macrolide resistance). There is commonality here with the synthetic ARGs previously described as contaminating laboratory reagents^8^. Co-occurrences with some of the above ARGs, in particular *blaCTXAM,* are hypothesised to be due to assembly artefacts, generated by the erroneous linkage of a synthetic ARG carrying *blaTEM* to chromosomal location of *blaTEM* with a second ARG.

### Geographically localised synthetic vector1associated ARGs

To capture a broader set of synthetic vector-associated ARGs, the above search of the NCBI wgs database was re-applied with a second abundant ARG, aminoglycoside-3’ – phosphotransferase *aph(3),* used in place of *blaTEM.* Identified hits with *blaTEM* co-occurring were removed. This search identified a comparatively limited set of organisms to which synthetic vector-associated *aph(3’)* was attributed (Supplementary Table 2), although it was able to identify synthetic vectors in pathogens outside of the *blaTEM-based* search, such as *Streptococcus pyogenes* (Supplementary Table 1). Identified hits include biological samples sequenced between 2009-2018, with these appearing to be geographically localised – the majority of identified species having been sequenced in China (54%), and others sequenced in India (15%), USA (15%), Germany (11%), Chile (2%) and Finland(2%). Patterns in regional and temporal appearances of synthetic vector associated ARGs can be observed in the above *blaTEM* set as well, although these are less pronounced (Supplementary Table 1).

### Synthetic vector-associated ARGs as reagent contaminants

Together, the observed breadth of phylogenetic groups in which these synthetic vector associated ARGs appear, the commonality to synthetic ARGs previously observed as contaminating, and the geographic and temporal distributions of synthetic vector associated ARGs, suggest ongoing and widespread reagent contamination. As mentioned, commonly used laboratory reagents have been a source of synthetic ARG contamination in the past, namely thermostable *Taq* polymerases^6,9^. In this work, possible sources of contamination could appear as encoded proteins on the identified synthetic vectors. As such, reagents appearing as encoded proteins and therefore contaminating were explored, and were shown to include thermostable DNA polymerase (as appearing on PRLD01000031), RNAse I (as appearing on JNCU01000114) and His^tagged Polynucleotide Kinase (as appearing on PIHF01000021). In the case of the identified polynucleotide kinase encoding synthetic vector, contamination appears to be geographically localised to China (Supplementary Table 2). The above reagents are used at various points in sequence generation^13,14^, and contamination of these and potentially other reagents with synthetic ARGs poses a challenge for the application of next-generation sequencing in clinical microbiology, potentially confounding results and impacting on patient management^15^.

### Literature examples of synthetic ARG contamination

In light of these findings, previous publications that highlight the appearance of ARGs in unexpected taxa should be reinterrogated. As an example, the finding of beta-lactamases in *Streptococcus agalactiae* and *Streptococcus uberis* at high frequencies (>90%) in Canada^16^, alongside high frequencies of aminoglycoside resistance (100%), can be explained by the presence of a synthetic vector carrying *blaTEM* and *aph(3’*) in all sequenced samples (Supplementary Figure 1). As a second example, the unexpected abundance of beta-lactamase expression in oceanic meta-transcriptomes^17^, including sea sediments^18^, open ocean derived sea phytoplankton^19^ and marine sea sponges^17^ can also be attributed to the presence of synthetic vectors carrying *blaTEM* (Supplementary Figure 2). In the meta-transcriptome of open ocean derived sea phytoplankton, *blaTEM* co-locates with an encoded lacZalpha peptide. In the meta-transcriptome of a sea sponge, *blaTEM* co-locates with a pUC-family origin of replication, and in the meta-transcriptomes of sea sediments collected at various depths, *blaTEM* appears to collocate with various synthetic vector sequences (Supplementary Figure 2). These data illustrate varying levels of contamination across samples, with some providing sufficient sequence coverage to recreate entire synthetic vectors and/or ARGs, and others providing partial coverage (Supplementary Figures 1 and 2). These findings should be considered when evaluating the abundance of ARGs in nature.

### Implications for diagnostic microbiology

Above we describe the appearance of synthetic vector-associated ARGs in next-generation sequences attributed to over 120 microbial species, including high priority pathogens^12^ (Table 1). Evidence presented here supports the hypothesis that this appearance is due to the ongoing and widespread contamination of reagents used in generating next-generation sequences. As mentioned, this has direct relevance for the application of next-generation sequencing in clinical microbiology for the diagnosis of antimicrobial resistance. In particular, this has the potential to generate inaccurate microbial genotypic information, potentially impacting patient management and outcomes^15^. In an effort to address the possibility of synthetic vector-associated ARG contamination, one could seek to modify the synthetic ARGs used in molecular biology so that they no longer reflect those found in nature. Other routes could include additional quality checks during the manufacture of reagents, especially for those employed in clinical microbiology^20^, or in processing next-generation sequence data.

## Conclusions

Here, to the best of our knowledge, we report the first description of the appearance of synthetic vector-associated ARGs as contaminating sequences across a broad phylogenetic space, as shown in varied and contemporary next-generation sequence-based data sets and assemblies. This finding has implications for the study of antibiotic resistance in clinical and environmental settings, and as next-generation sequencing becomes increasingly applied in clinical microbiology, could also impact clinical decision making in the near-future.

### Funding

The study was supported by internal funding.

### Transparency Declarations

None to declare

### Supplementary Data

Table S1, S2 and Figures S1 and S2 are available as Supplementary data.

---

**Supplementary Figure 1.**
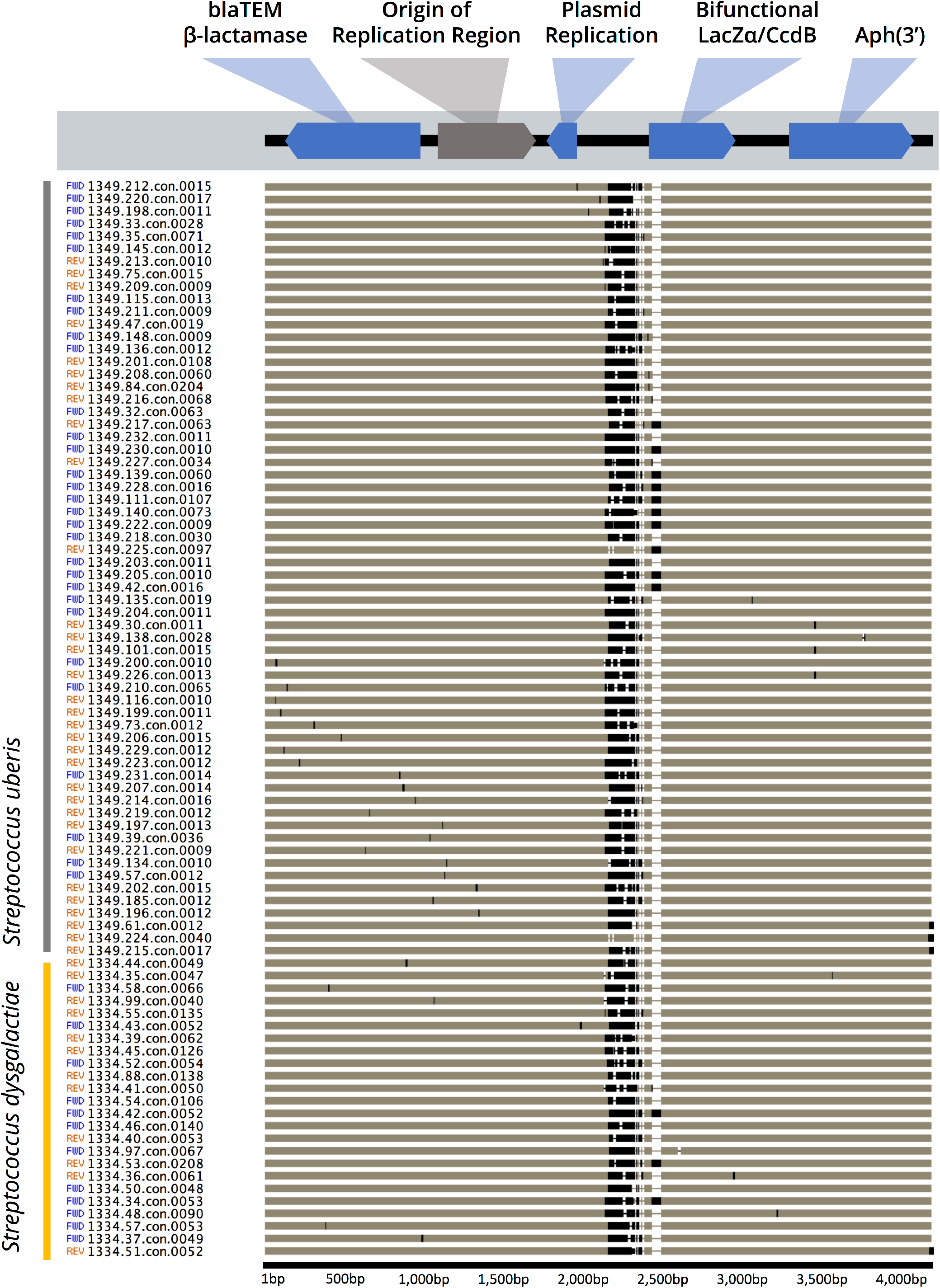
Alignment of synthetic vectors identified in Canadian *Streptococcus uberis* and *Streptococcus agalactiae* isolates sequenced by Vélez et al^18^

**Supplementary Figure 2.**
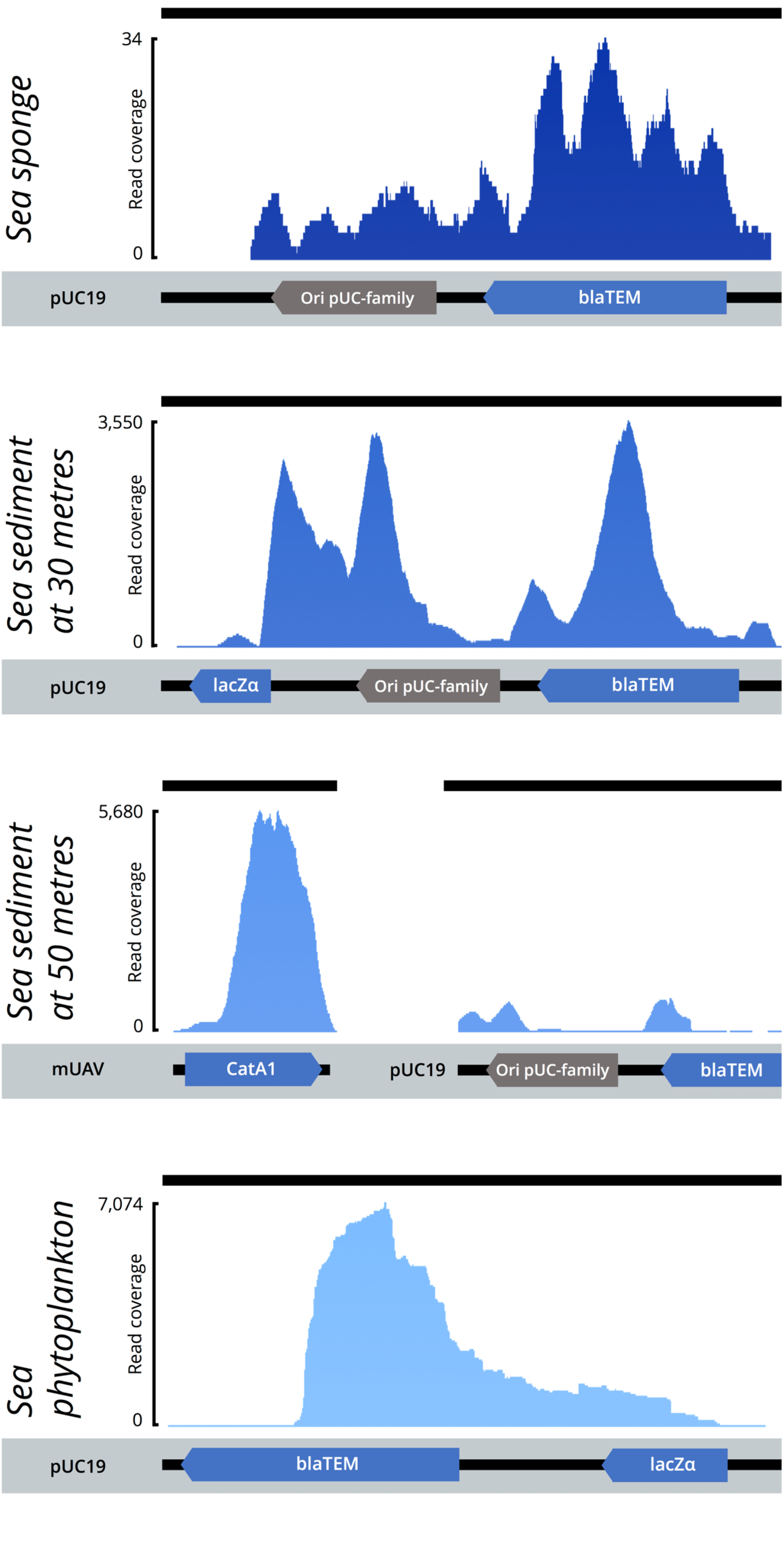
Coverage plot of oceanic metaCtranscriptomes mapped to synthetic vectors

**Supplementary Table 1.**
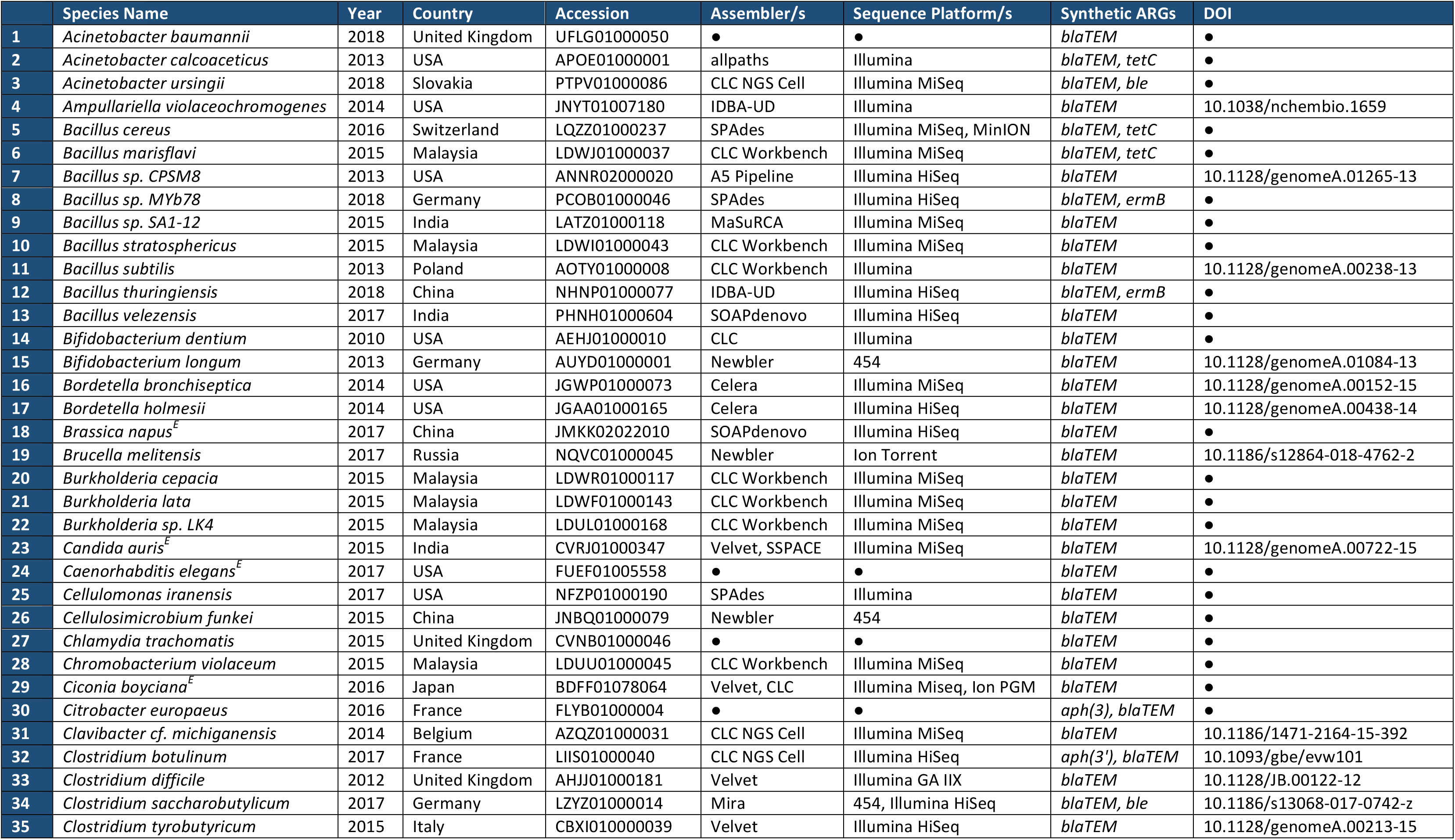
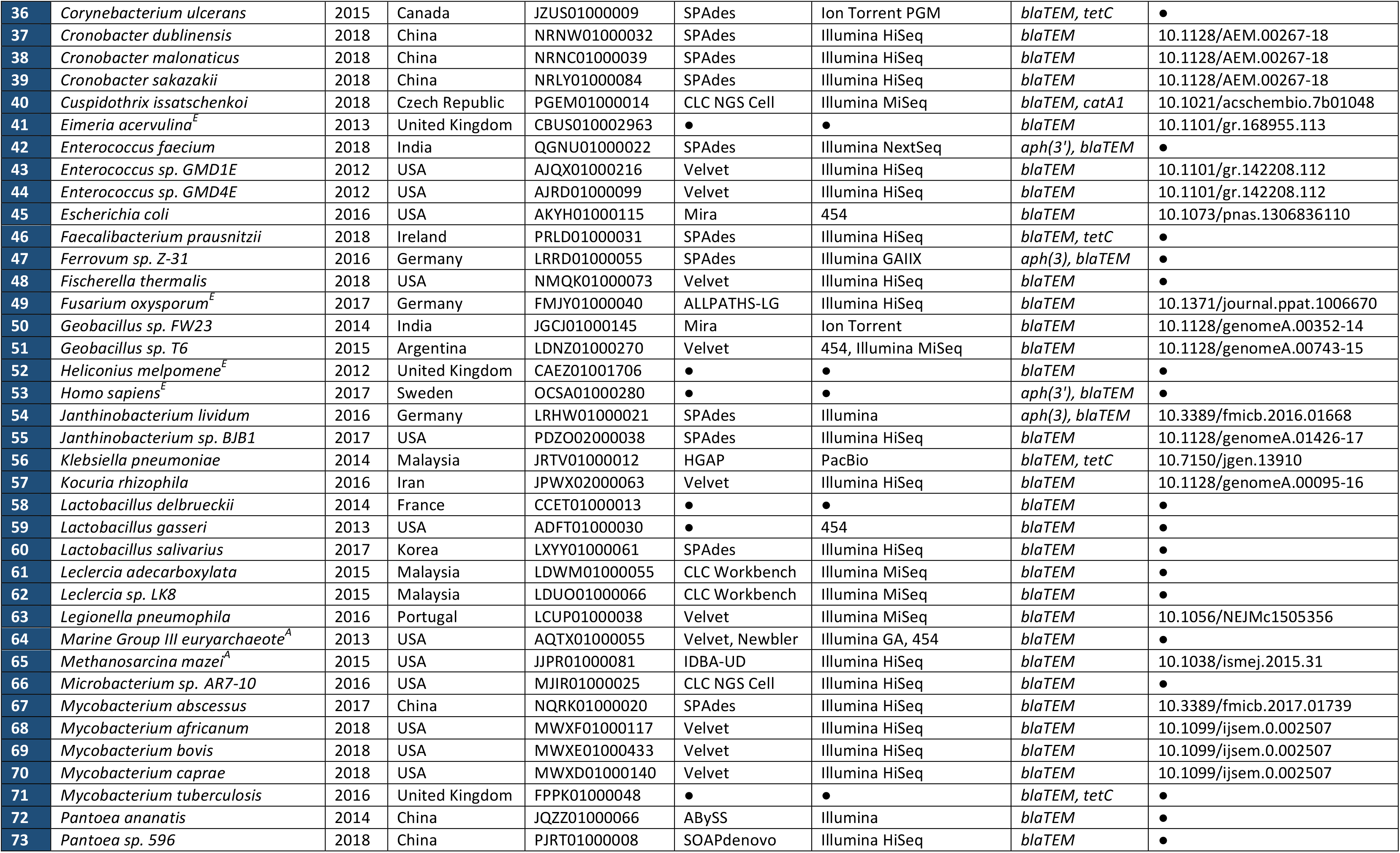
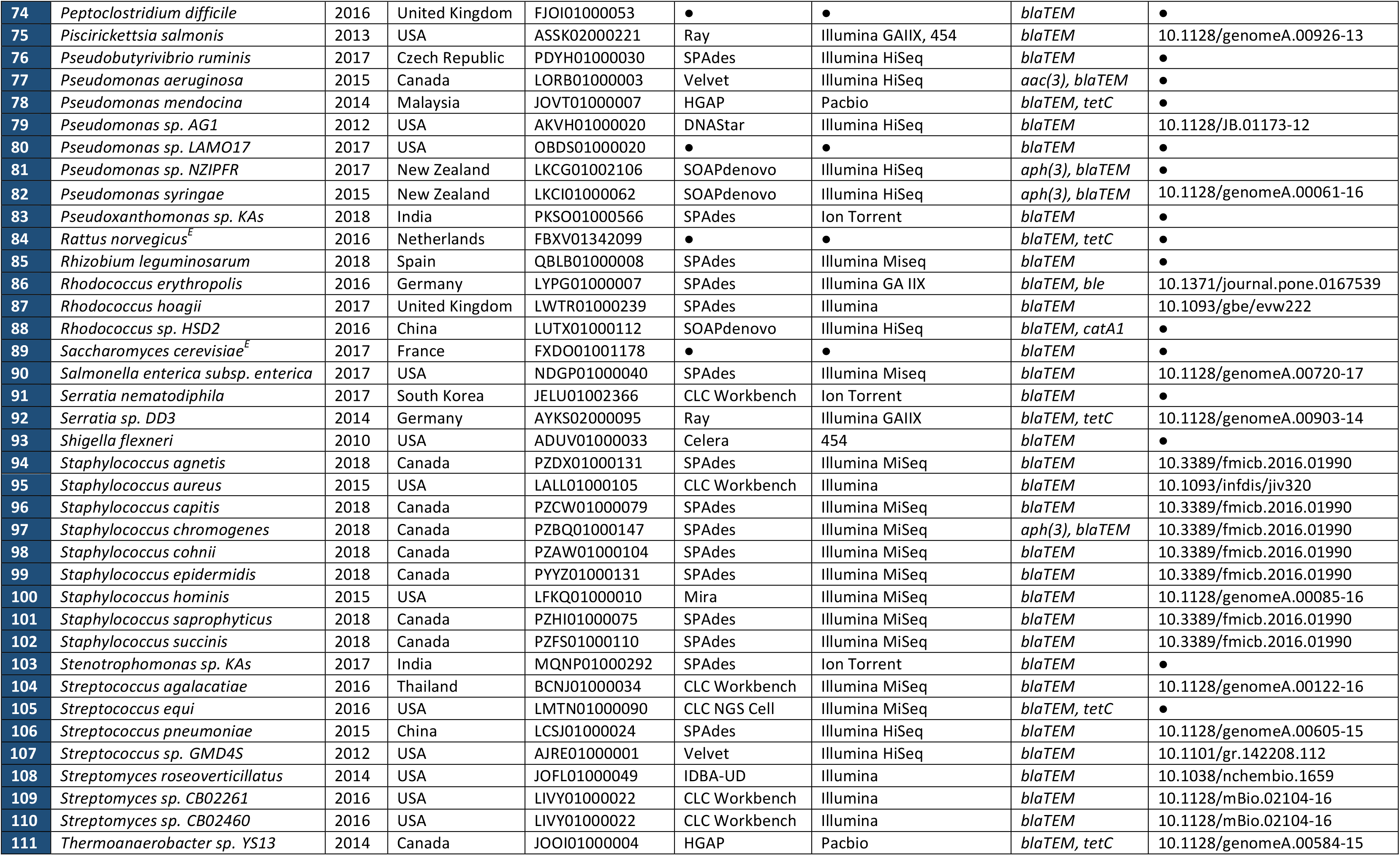
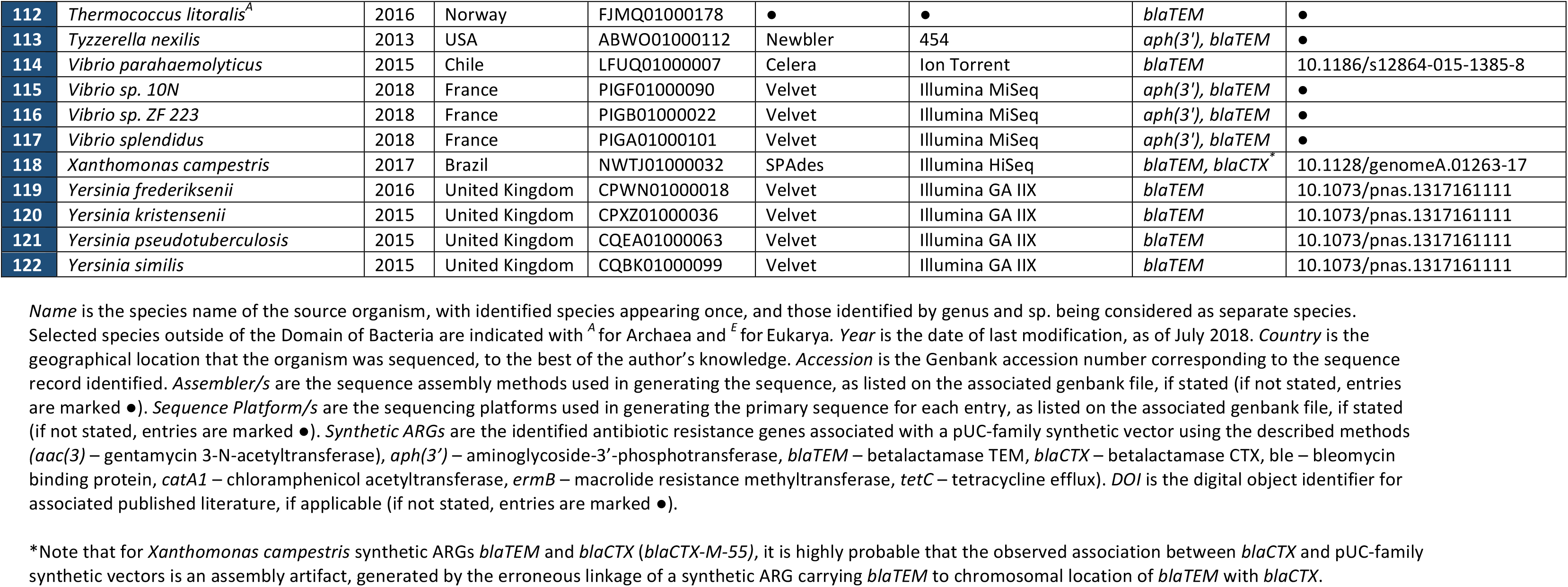
Examples of pUC-family synthetic vector contamination in the NCBI wgs database

**Supplementary Table 2.**
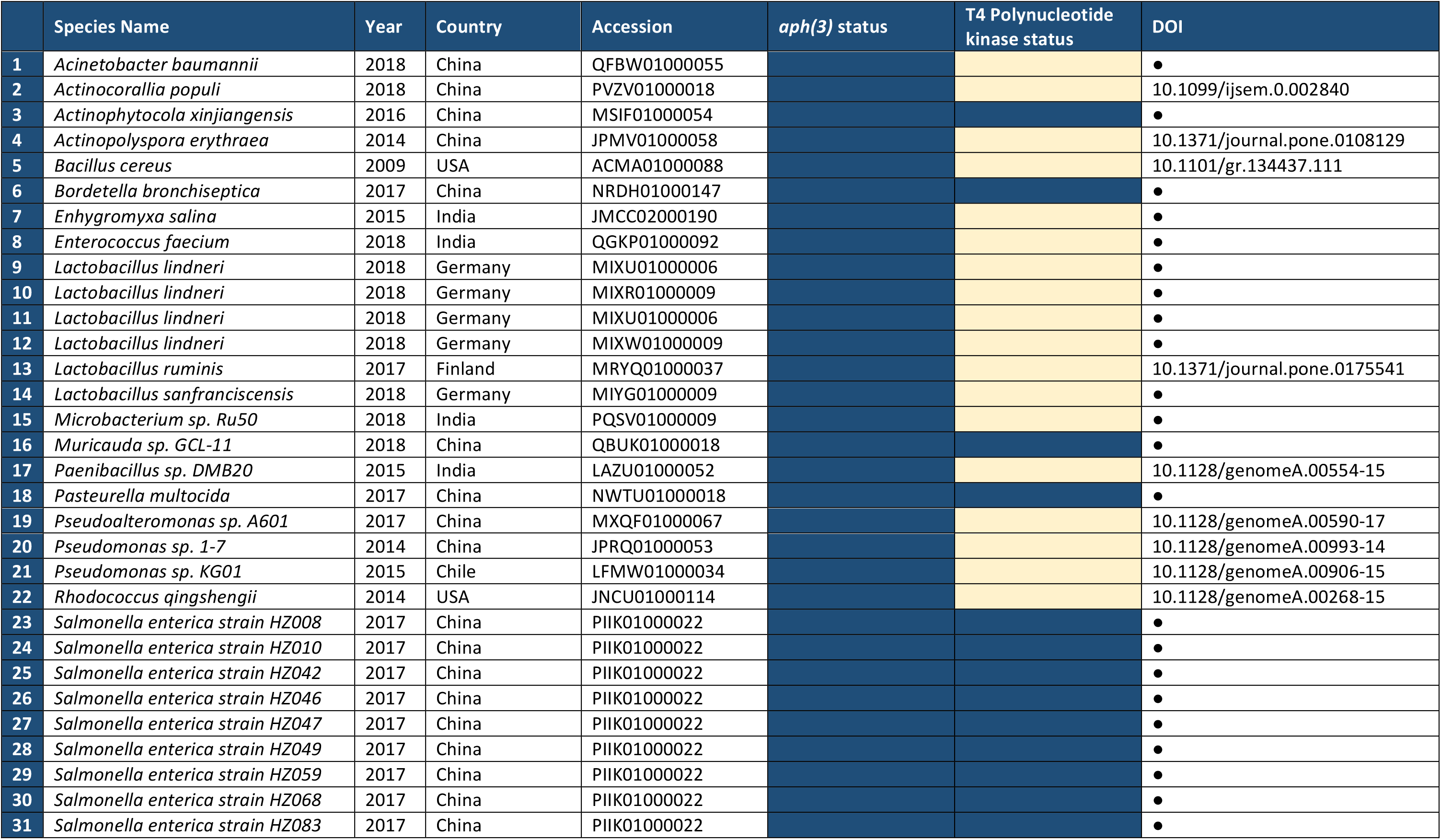
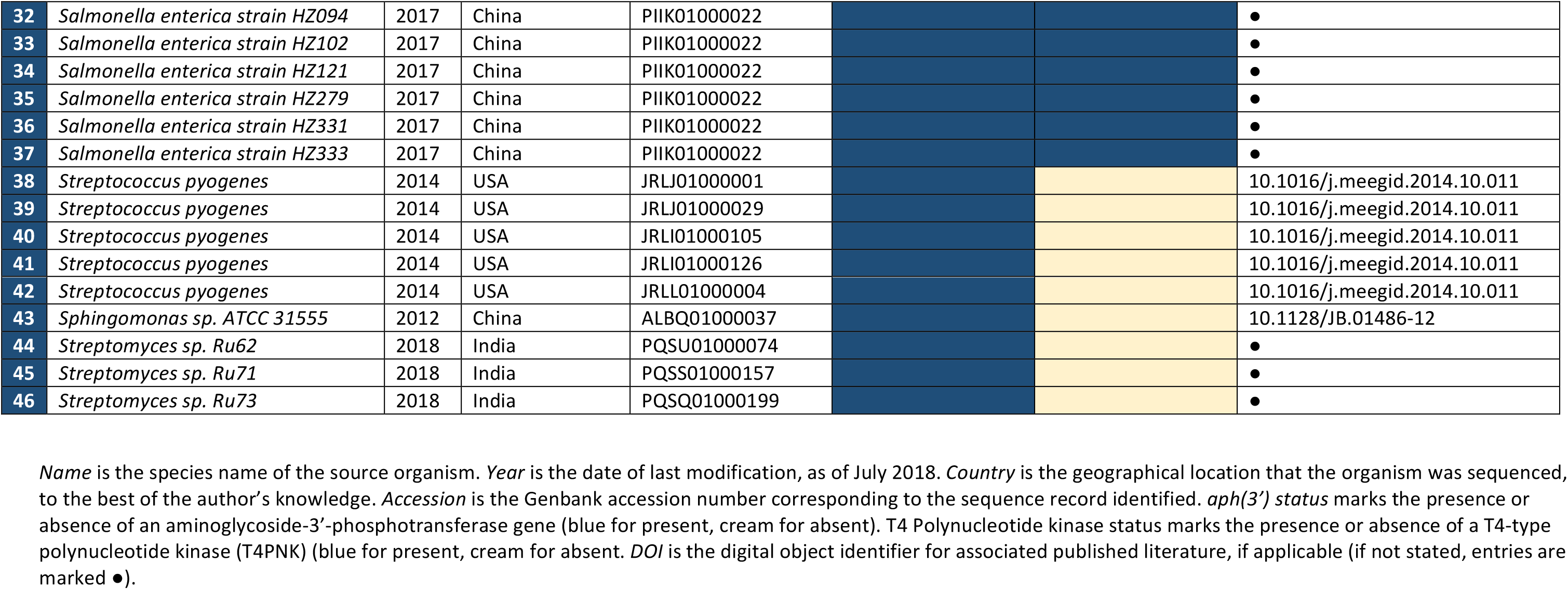
An example of geographically localized synthetic vector contamination

## Supplementary Methods

Appearance of synthetic antibiotic resistance gene (ARG) sequences in next-generation sequence databases was initially explored by searching for synthetic vectors of the pUC-family (Veieira and Messing 1982, Casali and Preston 2003). These were selected because:

i. pUC family vectors commonly carry the ARG *blaTEM* (Veieira and Messing 1982), encoding a betalactamase (Jacoby and Bush 2016), one of the most commonly used ARGCbased selectable markers in molecular biotechnology (Song *et al.* 2006)
ii. The ARG *blaTEM* described above is not typically found naturally in Gram-positive bacteria (Koncan *et al.* 2007), providing an early indication of synthetic vector contamination that could be explored if the *blaTEM* were to appear outside of the natural distribution in Gram-negative bacteria.
iii. pUC family vectors have been in use for decades throughout molecular biology and biotechnology, and are still used widely (Veieira and Messing 1982, Rodriguez 1988).
iv. Synthetic vectors of the pUC-family have a variable ARG repertoire allowing for selection with antibiotic exposures other than beta-lactams. Such pUC-family associated ARGs include those associated with aminoglycoside resistance (Pridmore 1987, Jeffrey and Joachim 1991), tetracycline resistance (Quigley and Reeves 1987), chloramphenicol resistance (Chang *et al.* 2015), bleomycin resistance (Austin *et al.* 1990) and others. There is therefore a breadth of possible synthetic ARGs that could be identified associated with contaminating pUC-family plasmids in next-generation sequence data.
v. This family of synthetic vectors has been hypothesized to be a source of contamination previously, with synthetic *blaTEM* being found in various laboratories as a contaminant in reagents used for diagnostic PCR (Song *et al.* 2006, Koncan *et al.* 2007, Chiang *et al.* 2005, Perron et al. 2006, Furlan *et al.* 2017).

Notable sequences commonly located on pUC-family synthetic vectors include the above *blaTEM* ARG, originating in RSF2124 and derived directly from pBR322 (Bolivar *et al.* 1977a, Bolivar *et al.* 1977b). This *blaTEM* carries various point mutations in derived synthetic vectors, generating amino acid changes *(blaTEM-1* as in pBR322, *blaTEM-116* having V84I and A184V as in pUC19 (Norrander *et al.* 1983), *blaTEM-171* having V84I as in pUC5 (Veieira and Messing 1982), *blaTEM-181* with A184V as in various pBAD plasmids (Zhang *et al.* 2017), and others (Zeil *et al.* 2016)). These vectors also commonly include a pMB1-type origin of replication region (Veieira and Messing 1982). These two features are shared with other synthetic vectors, such as pBR322 and related plasmids (Bolivar *et al.* 1977a, Bolivar *et al.* 1977b), as the original pUC series was derived from pBR322. Features of the pUC-family of synthetic vectors have also been included in modular plasmid systems, with these appearing in diverse plasmids (Charpenteir *et al.* 2004), as well as major modifications to pUCCfamily plasmids now being in use (Benes *et al.* 1993, Parke 1990). Aiming initially to identify synthetic ARGs by identifying the pUCCfamily *blaTEM* and associated sequences such as the origin of replication region could potentially capture varied synthetic vectors, inclusive of many points of contamination.

Areas chosen as markers of synthetic vectors in this work include the pUC-family *blaTEM* betalactamase (specifically *blaTEM-116* as found on pUC19 (Norrander *et al.* 1983) – Accession number L09137.2, nucleotide position 1626-2486bp) and origin of replication region (as found on pUC19 c Accession number L09137.2, nucleotide position 867-1455bp). As it has been claimed that *blaTEM-116* appears in nature, it cannot be considered as a synthetic ARG without being placed in a surrounding synthetic genetic context (Zeil *et al.* 2016, Jacoby and Bush 2016). The natural appearance of this ARG remains contentious however.

The characteristic sequences were entered as queries in BLASTn searches of two selected databases: (i) the NCBI (National Centre for Biotechnology and Information) nucleotide collection, consisting of GenBank (Benson *et al.* 2012), EMBL (Kanz *et al.* 2005), DDBJ (Mashima *et al.* 2016), PDB (Berman *et al.* 2000) and RefSeq (O’Leary *et al.* 2016) sequences but excluding expressed sequence tags (ESTs), (sequence-tagged sites (STSs), genome survey sequences (GSSs), whole-genome shotgun (WGS) sequences, transcriptome shotgun assembly (TSA) sequences, the Short Read Archive (SRA) and patent sequences, and (ii) a second search of the NCBI whole genome shotgun contigs database (wgs). Each of the two NCBI databases were chosen as they include many sequences for a breadth of organisms (Prokaryota, Eukaryota, Archaea), and are typically curated to include relevant metadata such as the sequence platform/s and assembly method/s used. Entries in these databases are also generally assembled contigs as well as draft or completed genomes, rather than short reads. This allows for the genetic context of an ARG to be explored, allowing for discrimination between naturally occurring and synthetic ARGs.

BLASTn searches were completed using NCBI BLAST online (Johnson *et al.* 2008) with the ‘megablast’ program. The search set was varied between open searches (nucleotide collection) and those directed towards certain taxa (wgs).

Criteria for a hit include

i. A BLAST aligned length of *blaTEM* (≥500 base pairs, ≥99.5% sequence identity across all of the *blaTEM* sequence aligned) and a length of synthetic vector sequence (≥100 base pairs, ≥99.5% sequence identity to one or more known synthetic vectors). As the majority of hits included a complete *blaTEM,* and averaged >4,000bp in length, a commonly identified synthetic vector structure was the associated origin of replication region, typically located 170bp downstream from the *blaTEM* gene.
ii. The above hits to *blaTEM* and a synthetic vector must be continuous, without gaps (Ns) or identified transposable/repeat element-associated sequences between the two areas
iii. Identified hits described as vector-containing, or otherwise transgenic, were excluded as synthetic vectors are likely not contaminating in these instances.

Comparison and alignment to known synthetic vectors was carried out through BLASTn searches of the nucleotide database on NCBI BLAST online, and using various alignment and mapping tools in Geneious 8.1.9 (Kearse et al. 2012) including MUSCLE (Edgar 2004), progressiveMAUVE (Darling *et al.* 2010), and Geneious read mapper. Associated synthetic ARGs to *blaTEM* were identified using a combination of:

i. Open reading frame detection in Geneious 8.1.9, and BLASTp of these towards NCBI non-redundant protein sequences with NCBI BLAST online
ii. ABRicate-based detection of ARGs, followed by manual curation (https://github.com/tseemann/abricate)

Identified hits were ordered in a table, including information garnered from Genbank files and associated publications for each hit (species name, year last modified, country of origin, genbank accession number, assembly method/s used, sequence platform/s used, DOI of associated publication, title).

The appearance of synthetic ARG contamination across broad phylogenetic space was illustrated through the annotation of a simplified phylogenetic tree of life, as appearing on the Interactive Tree of Life (iTOL v4)(Letunic and Bork 2016). The simplified tree shows the Archaea, Eukarya and the Firmicute branch of the Prokarya as they appear on iTOL v4. These branches were selected as *blaTEM* appears in Gram-negatives naturally, while the appearance of *blaTEM* in organisms belonging to the shown groups is atypical. Selected examples of Archaea and Eukarya with synthetic vector-associated ARGs are shown on this phylogenetic tree, and in Supplementary Table 1.

The above search was repeated with the second most abundant ARG, aminoglycoside-3’-phosphotransferase *aph(3)*, used in place of *blaTEM.* Identified hits with *blaTEM* cooccurring were removed, as they appear in the initial search.

Criteria for a hit include:

i. A BLAST aligned length of *aph(3)* (≥500 base pairs, ≥99.5% sequence identity across all of the *aph(3)* sequence aligned) and a length of synthetic vector sequence (≥100 base pairs, ≥99.5% sequence identity to one or more known synthetic vectors).
ii. The above hits to *aph(3)* and a synthetic vector must be continuous, without gaps (Ns) or identified transposable/repeat element-associated sequences between the two areas
iii. Identified hits described as vector-containing, or otherwise transgenic, were excluded as synthetic vectors are likely not contaminating in these instances.

Two versions of *aph(3)* were applied in the above search, as two distinct synthetic vector-associated *aph(3)* genes were identified, having approximately 33% sequence identity on a protein level. Sequences of these two versions used in the search of the NCBI wgs database were those appearing on MIXW01000009 (encoded on nucleotide positions 988-1,803) and on LIIS01000040 (encoded on nucleotide positions 2,113-2,907), for each aph(3) version respectively. The *aph(3)* version appearing on LIIS01000040 was generally associated with *blaTEM,* and so hits removed from this analysis, while the *aph(3)* appearing on MIXW01000009 was not associated with *blaTEM* and makes up the entirety of hits shown in Supplementary Table 2.

Synthetic sequences encoding laboratory reagents were identified through manual curation of large open reading frames (over 600bp) in Geneious 8.1.9, and BLASTp of these open reading frames towards NCBI non-redundant protein sequences with NCBI BLAST online.

Selected literature examples of specific ARGs appearing in unexpected taxa were hypothesised to be due to synthetic vector contamination. These include:

i. Appearance of *blaTEM* in *Streptococcus agalactiae* and *Streptococcus uberis* at high frequencies (>90%) in Canada (Vélez et al. 2017), alongside aminoglycoside resistance. This is unexpected, with the study authors stating *‘The presence of encoding regions for β-lactamases in this population of isolates, which are catalase-negative Gram-positive cocci, is a novel finding’.*
ii. Abundance of blaTEM expression in oceanic meta-transcriptomes (Versluis *et al.* 2015), including sea sediments (Orsi *et al.* 2013), open ocean derived sea phytoplankton (Gifford *et al.* 2011) and marine sponges (Versluis *et al.* 2015). This is unexpected, with the study authors stating *‘Surprisingly, in sea bacterioplankton, the expression of AR genes was more than one order of magnitude higher than in any other ecological niche. It was comprised of two highly similar β-lactamase genes, namely blaTEM-1 and blaTEM-116… The blaTEM-1 gene is commonly present in cloning vectors, and therefore lab contamination can be suspected, however, this possibility is disputed by the fact that the gene is detected in two separate marine metatranscriptome datasets that have been independently generated. The functional role of a β-lactamase in a marine environment is yet unknown.’* (Versluis *et al.* 2015).

The hypothesis that these unexpected appearances of ARGs was due to synthetic vector contamination was explored through analysis of both primary reads and assemblies. A focus was placed on determining the genetic context of identified ARGs to determine if they were of synthetic origins. Assembled contigs attributed to Canadian *Streptococcus agalactiae* and *Streptococcus uberis* were accessed from the Pathosystems Resource Integration Center (PATRIC)(Wattam *et al.* 2017), with contigs carrying *blaTEM* being identified using Geneious Read Mapper *(blaTEM* provided as a reference sequence as it appears in Accession number L09137.2, nucleotide position 1626-2486bp), before being aligned using MUSCLE, and annotated using BLASTp of identified open reading frames in NCBI BLAST online and Geneious 8.1.9.

In addressing the abundance of *blaTEM* expression in oceanic meta-transcriptomes, primary reads for the various oceanic meta-transcriptomes mentioned in the study were accessed from the European Nucleotide Archive (ENA)(Leinonen et al. 2011). These were sample accessions/Bioproject numbers CAM_PROJ_Sapelo2008 (open ocean, Gifford *et al.* 2011), ERS566232 (sea sponge, Versluis *et al.* 2015) and SRR571458, SRR948190 and SRR948292 (sea sediments, Orsi *et al.* 2013). Individual datasets were mapped to pUC19 (Accession number L09137.2) using Geneious read mapper, simultaneously identifying reads mapping to *blaTEM* and those mapping to other areas of pUC-family vectors as described above. Settings for Geneious read mapping here were a custom sensitivity, allowing gaps (maximum of 15%), mismatches (maximum of 20%) and ambiguities (maximum 16). An index word length of 11, and a word length of 12 were applied, along with the *‘Accurately map reads to repeat regions’* and *‘Map multiple best matches to all’* settings. Meta-transcriptomic datasets were also mapped to synthetic ARGs identified earlier in this work in NCBI databases *(aac(3), aph(3’), ble, catAl, ermB, tetC).* This was to illustrate how in some settings, contamination with synthetic ARGs leads to apparent expression of diverse ARGs other than *blaTEM.* Coverage plots were exported from Geneious 8.1.9. Note that in the case of the open ocean and sea sponge meta-transcriptomes, coverage is continuous across the areas shown. In the sea sediment samples, coverage is discontinuous between the pUC-origin of replication region and the *blaTEM* gene, potentially due to additional gene content at these sites in the identified synthetic vectors.

## References

1. O’Neill, J. (2014). Antimicrobial Resistance: Tackling a Crisis for the Health and Wealth of Nations. The Review on Antimicrobial Resistance. (https://amr-review.org/sites/default/files/160525_Final%20paper_with%20cover.pdf)

2. Deurenberg, R. H., Bathoorn, E., Chlebowicz, M. A., Couto, N., Ferdous, M., García-Cobos, S., … Rossen, J. W. A. (2017). Application of next generation sequencing in clinical microbiology and infection prevention. Journal of Biotechnology, 243, 16–24. https://doi.org/10.1016/j.jbiotec.2016.12.022

3. Ellington, M. J., Ekelund, O., Aarestrup, F. M., Canton, R., Doumith, M., Giske, C., … Woodford, N. (2017). The role of whole genome sequencing in antimicrobial susceptibility testing of bacteria: report from the EUCAST Subcommittee. Clinical Microbiology and Infection, 23(1), 2–22. https://doi.org/10.1016/j.cmi.2016.11.012

4. Otto, M. (2017). Next-generation sequencing to monitor the spread of antimicrobial resistance. Genome Medicine, 9(1), 68. https://doi.org/10.1186/s13073-017-0461-x

5. Jiang, X., Ellabaan, M. M. H., Charusanti, P., Munck, C., Blin, K., Tong, Y., … Lee, S. Y. (2017). Dissemination of antibiotic resistance genes from antibiotic producers to pathogens. Nature Communications, 8, 15784. https://doi.org/10.1038/ncomms15784

6. Koncan, R., Valverde, A., Morosini, M.-I., García-Castillo, M., Cantón, R., Cornaglia, G., … del Campo, R. (2007). Learning from mistakes: Taq polymerase contaminated with (β-lactamase sequences results in false emergence of Streptococcus pneumoniae containing TEM. Journal of Antimicrobial Chemotherapy, 60(3), 702–703. https://doi.org/10.1093/jac/dkm239

7. Chiang, C., Liu, C., Weng, L., Wang, N., and Liaw, G. (2005). Presence of beta-lactamase gene TEM-1 DNA sequence in commercial Taq DNA polymerase. Journal of Clinical Microbiology, 43(1), 530–1. https://doi.org/10.1128/JCM.43.1.530-531.2005

8. Perron, A., Raymond, P., Simard, R. (2006). The Occurrence of Antibiotic Resistance Genes in Taq Polymerases and a Decontamination Method Applied to the Detection of Genetically Modified Crops. Biotechnology Letters, 28(5), 321–325. https://doi.org/10.1007/s10529-005-5931-3

9. Furlan, J. P. R., Stehling, E. G., Pitondo-Silva, A. (2017). Importance of Sequencing To Determine Functional bla TEM Variants. Antimicrobial Agents and Chemotherapy, 61(5), e00237–17. https://doi.org/10.1128/AAC.00237-17

10. Yang, S., Rothman, R. E. (2004). PCR-based diagnostics for infectious diseases: uses, limitations, and future applications in acute-care settings. The Lancet Infectious Diseases, 4(6), 337–348. https://doi.org/10.1016/S1473-3099(04)01044-8

11. Vieira, J., Messing, J. (1982). The pUC plasmids, an M13mp7-derived system for insertion mutagenesis and sequencing with synthetic universal primers. Gene, 19(3), 259–268. https://doi.org/10.1016/0378-1119(82)90015-4

12. Tacconelli, E., Carrara, E., Savoldi, A., Harbarth, S., Mendelson, M., Monnet, D. L., … Zorzet, A. (2018). Discovery, research, and development of new antibiotics: the WHO priority list of antibiotic-resistant bacteria and tuberculosis. The Lancet Infectious Diseases, 18(3), 318–327. https://doi.org/10.1016/S1473-3099(17)30753-3

13. Chen, C.-Y. (2014). DNA polymerases drive DNA sequencing-by-synthesis technologies: both past and present. Frontiers in Microbiology, 5. https://doi.org/10.3389/fmicb.2014.00305

14. Healey, A., Furtado, A., Cooper, T., Henry, R. J. (2014). Protocol: a simple method for extracting next-generation sequencing quality genomic DNA from recalcitrant plant species. Plant Methods, 10(1), 21. https://doi.org/10.1186/1746-4811-10-21

15. Motro, Y., Moran-Gilad, J. (2017). Next-generation sequencing applications in clinical bacteriology. Biomolecular Detection and Quantification, 14, 1–6. https://doi.org/10.1016/j.bdq.2017.10.002

16. Vélez, J. R., Cameron, M., Rodríguez-Lecompte, J. C., Xia, F., Heider, L. C., Saab, M., … Sánchez, J. (2017). Whole-Genome Sequence Analysis of Antimicrobial Resistance Genes in Streptococcus uberis and Streptococcus dysgalactiae Isolates from Canadian Dairy Herds. Frontiers in Veterinary Science, 4. https://doi.org/10.3389/fvets.2017.00063

17. Versluis, D., D’Andrea, M. M., Ramiro Garcia, J., Leimena, M. M., Hugenholtz, F., Zhang, J., … Passel, M. W. J. van. (2015). Mining microbial metatranscriptomes for expression of antibiotic resistance genes under natural conditions. Scientific Reports, 5(1), 11981. https://doi.org/10.1038/srep11981

18. Stewart, F. J., Sharma, A. K., Bryant, J. A., Eppley, J. M., DeLong, E. F. (2011). Community transcriptomics reveals universal patterns of protein sequence conservation in natural microbial communities. Genome Biology, 12(3), R26. https://doi.org/10.1186/gb-2011-12-3-r26

19. Gifford, S. M., Sharma, S., Rinta-Kanto, J. M., Moran, M. A. (2011). Quantitative analysis of a deeply sequenced marine microbial metatranscriptome. The ISME Journal, 5(3), 461–472. https://doi.org/10.1038/ismej.2010.141

20. Salter, S. J., Cox, M. J., Turek, E. M., Calus, S. T., Cookson, W. O., Moffatt, M. F., … Walker, A. W. (2014). Reagent and laboratory contamination can critically impact sequence-based microbiome analyses. BMC Biology, 12(1), 87. https://doi.org/10.1186/s12915-014-0087-z

## Supplementary References

Ahmad, N., Chong, T. M., Hashim, R., Shukor, S., Yin, W.-F., & Chan, K.-G. (2015). Draft Genome of Multidrug-Resistant Klebsiella pneumoniae 223/14 Carrying KPC-6, Isolated from a General Hospital in Malaysia. Journal of Genomics, 3, 97–98. https://doi.org/10.7150/jgen.13910

Alvarez, C., Kukutla, P., Jiang, J., Yu, W., & Xu, J. (2012). Draft Genome Sequence of Pseudomonas sp. Strain Ag1, Isolated from the Midgut of the Malaria Mosquito Anopheles gambiae. Journal of Bacteriology, 194(19), 5449–5449. https://doi.org/10.1128/JB.01173-12

Anastasi, E., MacArthur, I., Scortti, M., Alvarez, S., Giguère, S., & Vázquez-Boland, J. A. (2016). Pangenome and Phylogenomic Analysis of the Pathogenic Actinobacterium Rhodococcus equi. Genome Biology and Evolution, 8(10), 3140–3148. https://doi.org/10.1093/gbe/evw222

Areechon, N., Kannika, K., Hirono, I., Kondo, H., & Unajak, S. (2016). Draft Genome Sequences of Streptococcus agalactiae Serotype la and III Isolates from Tilapia Farms in Thailand. Genome Announcements, 4(2). https://doi.org/10.1128/genomeA.00122-16

Austin, B., Hall, R. M., & Tyler, B. M. (1990). Optimized vectors and selection for transformation of Neurospora crassa and Aspergillus nidulans to bleomycin and phleomycin resistance. Gene, 93(1), 157–162. https://doi.org/10.1016/0378-1119(90)90152-H

Benes, V., Hostomský, Z., Arnold, L., & Pačes, V. (1993). M13 and pUC vectors with new unique restriction sites for cloning. Gene, 130(1), 151–152. https://doi.org/10.1016/0378-1119(93)90360-F

Benson, D. A., Karsch-Mizrachi, I., Clark, K., Lipman, D. J., Ostell, J., & Sayers, E. W. (2012). GenBank. Nucleic Acids Research, 40(D1), D48–D53. https://doi.org/10.1093/nar/gkr1202

Berman, H. M. (2000). The Protein Data Bank. Nucleic Acids Research, 28(1), 235–242. https://doi.org/10.1093/nar/28.1.235

Bessen, D. E., McShan, W. M., Nguyen, S. V., Shetty, A., Agrawal, S., & Tettelin, H. (2015). Molecular epidemiology and genomics of group A Streptococcus. Infection, Genetics and Evolution, 33, 393–418. https://doi.org/10.1016/j.meegid.2014.10.011

Bettina, A. M., Doing, G., O’Brien, K., Perron, G. G., & Jude, B. A. (2018). Draft Genome Sequences of Phenotypically Distinct Janthinobacterium sp. Isolates Cultured from the Hudson Valley Watershed. Genome Announcements, 6(3). https://doi.org/10.1128/genomeA.01426-17

Bolivar, F., Rodriguez, R. L., Betlach, M. C., & Boyer, H. W. (1977). Construction and characterization of new cloning vehicles I. Ampicillin-resistant derivatives of the plasmid pMB9. Gene, 2(2), 75–93. https://doi.org/10.1016/0378-1119(77)90074-9

Bolivar, F., Rodriguez, R. L., Greene, P. J., Betlach, M. C., Heyneker, H. L., Boyer, H. W., … Falkow, S. (1977). Construction and characterization of new cloning vehicle. II. A multipurpose cloning system. Gene, 2(2), 95–113. https://doi.org/10.1016/0378-1119(77)90000-2

Brouwer, M. S. M., Allan, E., Mullany, P., & Roberts, A. P. (2012). Draft Genome Sequence of the Nontoxigenic Clostridium difficile Strain CD37. Journal of Bacteriology, 194(8), 2125–2126. https://doi.org/10.1128/JB.00122-12

Chang, B., Ray, A., Tsuei, T., Wan, R. (2015) Construction of an enlarged pUC19 vector with a rop gene designed to study plasmid maintenance in Escherichia coli. JEMI, 19, 1–5

Casali, N., & Preston, A. (2003). E. coli Plasmid Vectors (Vol. 235). New Jersey: Humana Press. https://doi.org/10.1385/1592594093

Charpentier, E., Anton, A. I., Barry, P., Alfonso, B., Fang, Y., & Novick, R. P. (2004). Novel Cassette-Based Shuttle Vector System for Gram-Positive Bacteria. Applied and Environmental Microbiology, 70(10), 6076–6085. https://doi.org/10.1128/AEM.70.10.6076-6085.2004

Chen, D., Feng, J., Huang, L., Zhang, Q., Wu, J., Zhu, X., … Xu, Z. (2014). Identification and Characterization of a New Erythromycin Biosynthetic Gene Cluster in Actinopolyspora erythraea YIM90600, a Novel Erythronolide-Producing Halophilic Actinomycete Isolated from Salt Field. PLoS ONE, 9(9), e108129. https://doi.org/10.1371/journal.pone.0108129

Chiang, C.-S., Liu, C.-P., Weng, L.-C., Wang, N.-Y., & Liaw, G.-J. (2005). Presence of – Lactamase Gene TEM-1 DNA Sequence in Commercial Taq DNA Polymerase. Journal of Clinical Microbiology, 43(1), 530–531. https://doi.org/10.1128/JCM.43.1.530-531.2005

Correia, A. M., Ferreira, J. S., Borges, V., Nunes, A., Gomes, B., Capucho, R., … Gomes, J. P. (2016). Probable Person-to-Person Transmission of Legionnaires’ Disease. New England Journal of Medicine, 374(5), 497–498. https://doi.org/10.1056/NEJMc1505356

Darling, A. E., Mau, B., & Perna, N. T. (2010). progressiveMauve: Multiple Genome Alignment with Gene Gain, Loss and Rearrangement. PLoS ONE, 5(6), e11147. https://doi.org/10.1371/journal.pone.0011147

Doroghazi, J. R., Albright, J. C., Goering, A. W., Ju, K.-S., Haines, R. R., Tchalukov, K. A., … Metcalf, W. W. (2014). A roadmap for natural product discovery based on large-scale genomics and metabolomics. Nature Chemical Biology, 10(11), 963–968. https://doi.org/10.1038/nchembio.1659

Edgar, R. C. (2004). MUSCLE: multiple sequence alignment with high accuracy and high throughput. Nucleic Acids Research, 32(5), 1792–1797. https://doi.org/10.1093/nar/gkh340

Eppinger, M., McNair, K., Zogaj, X., Dinsdale, E. A., Edwards, R. A., & Klose, K. E. (2013). Draft Genome Sequence of the Fish Pathogen Piscirickettsia salmonis. Genome Announcements, 1(6). https://doi.org/10.1128/genomeA.00926-13

Fitzsimons, M. S., Novotny, M., Lo, C.-C., Dichosa, A. E. K., Yee-Greenbaum, J. L., Snook, J. P., … Han, C. S. (2013). Nearly finished genomes produced using gel microdroplet culturing reveal substantial intraspecies genomic diversity within the human microbiome. Genome Research, 23(5), 878–888. https://doi.org/10.1101/gr.142208.112

Furlan, J. P. R., Stehling, E. G., & Pitondo-Silva, A. (2017). Importance of Sequencing To Determine Functional bla TEM Variants. Antimicrobial Agents and Chemotherapy, 61(5). https://doi.org/10.1128/AAC.00237-17

Gifford, S. M., Sharma, S., Rinta-Kanto, J. M., & Moran, M. A. (2011). Quantitative analysis of a deeply sequenced marine microbial metatranscriptome. The ISME Journal, 5(3), 461–472. https://doi.org/10.1038/ismej.2010.141

Haack, F. S., Poehlein, A., Kröger, C., Voigt, C. A., Piepenbring, M., Bode, H. B., … Streit, W. R. (2016). Molecular Keys to the Janthinobacterium and Duganella spp. Interaction with the Plant Pathogen Fusarium graminearum. Frontiers in Microbiology, 7. https://doi.org/10.3389/fmicb.2016.01668

Harvill, E. T., Goodfield, L. L., Ivanov, Y., Smallridge, W. E., Meyer, J. A., Cassiday, P. K., … Losada, L. (2014). Genome Sequences of Nine Bordetella holmesii Strains Isolated in the United States. Genome Announcements, 2(3). https://doi.org/10.1128/genomeA.00438-14

Hazen, T. H., Sahl, J. W., Fraser, C. M., Donnenberg, M. S., Scheutz, F., & Rasko, D. A. (2013). Refining the pathovar paradigm via phylogenomics of the attaching and effacing Escherichia coli. Proceedings of the National Academy of Sciences, 110(31), 12810–12815. https://doi.org/10.1073/pnas.1306836110

Hughes, G. L., Raygoza Garay, J. A., Koundal, V., Rasgon, J. L., & Mwangi, M. M. (2016). Genome Sequences of Staphylococcus hominis Strains ShAs1, ShAs2, and ShAs3, Isolated from the Asian Malaria Mosquito Anopheles stephensi. Genome Announcements, 4(2). https://doi.org/10.1128/genomeA.00085-16

Ip, M., Ma, H., Li, C., Tsui, S., & Zhou, H. (2015). Draft Genome Sequences of Two Streptococcus pneumoniae Serotype 19F Sequence Type 271 Clinical Isolates with Low- and High-Level Cefotaxime Resistance. Genome Announcements, 3(3). https://doi.org/10.1128/genomeA.00605-15

Jacoby, G. A., & Bush, K. (2016). The Curious Case of TEM-116. Antimicrobial Agents and Chemotherapy, 60(11), 7000–7000. https://doi.org/10.1128/AAC.01777-16

Jeffrey, V., & Joachim, M. (1991). New pUC-derived cloning vectors with different selectable markers and DNA replication origins. Gene, 100, 189–194. https://doi.org/10.1016/0378-1119(91)90365-1

Johnson, M., Zaretskaya, I., Raytselis, Y., Merezhuk, Y., McGinnis, S., & Madden, T. L. (2008). NCBI BLAST: a better web interface. Nucleic Acids Research, 36(Web Server), W5–W9. https://doi.org/10.1093/nar/gkn201

Kant, R., Palva, A., & von Ossowski, I. (2017). An in silico pan-genomic probe for the molecular traits behind Lactobacillus ruminis gut autochthony. PLOS ONE, 12(4), e0175541. https://doi.org/10.1371/journal.pone.0175541

Kanz, C. (2004). The EMBL Nucleotide Sequence Database. Nucleic Acids Research, 33(Database issue), D29–D33. https://doi.org/10.1093/nar/gki098

Kazmierczak, R. A., Best, A. A., Nguyen, D., & Eisenstark, A. (2017). Whole-Genome Shotgun Sequences of Salmonella enterica Serovar Typhimurium Lilleengen Type Strains LT1, LT18, LT19, LT20, LT21, and LT22. Genome Announcements, 5(30). https://doi.org/10.1128/genomeA.00720-17

Kearse, M., Moir, R., Wilson, A., Stones-Havas, S., Cheung, M., Sturrock, S., … Drummond, A. (2012). Geneious Basic: An integrated and extendable desktop software platform for the organization and analysis of sequence data. Bioinformatics, 28(12), 1647–1649. https://doi.org/10.1093/bioinformatics/btsl99

Khairy, H., Meinert, C., Wübbeler, J. H., Poehlein, A., Daniel, R., Voigt, B., … Steinbüchel, A. (2016). Genome and Proteome Analysis of Rhodococcus erythropolis MI2: Elucidation of the 4,4ʹ-Dithiodibutyric Acid Catabolism. PLOS ONE, 11(12), e0167539. https://doi.org/10.1371/journal.pone.0167539

Koncan, R., Valverde, A., Morosini, M.-l., García-Castillo, M., Cantón, R., Cornaglia, G., … del Campo, R. (2007). Learning from mistakes: Taq polymerase contaminated with β-lactamase sequences results in false emergence of Streptococcus pneumoniae containing TEM. Journal of Antimicrobial Chemotherapy, 60(3), 702–703. https://doi.org/10.1093/jac/dkm239

Krzyzanowska, D. M., Iwanicki, A., Ossowicki, A., Obuchowski, M., & Jafra, S. (2013). Genome Sequence of Bacillus subtilis MB73/2, a Soil Isolate Inhibiting the Growth of Plant Pathogens Dickeya spp. and Rhizoctonia solani. Genome Announcements, 1(3). https://doi.org/10.1128/genomeA.00238-13

Kust, A., Mareš, J., Jokela, J., Urajová, P., Hájek, J., Saurav, K., … Hrouzek, P. (2018). Discovery of a Pederin Family Compound in a Nonsymbiotic Bloom-Forming Cyanobacterium. ACS Chemical Biology, 13(5), 1123–1129. https://doi.org/10.1021/acschembio.7b01048

Leinonen, R., Akhtar, R., Birney, E., Bower, L., Cerdeno-Tarraga, A., Cheng, Y., … Cochrane, G. (2011). The European Nucleotide Archive. Nucleic Acids Research, 39(Database), D28–D31. https://doi.org/10.1093/nar/gkq967

Letunic, I., & Bork, P. (2016). Interactive tree of life (iTOL) v3: an online tool for the display and annotation of phylogenetic and other trees. Nucleic Acids Research, 44(W1), W242–W245. https://doi.org/10.1093/nar/gkw290

Li, B., Yang, S., Chu, H., Zhang, Z., Liu, W., Luo, L., … Xu, X. (2017). Relationship between Antibiotic Susceptibility and Genotype in Mycobacterium abscessus Clinical Isolates. Frontiers in Microbiology, 8. https://doi.org/10.3389/fmicb.2017.01739

Li, J., Cheng, Y., Wang, D., Li, J., Wang, Y., Han, W., & Li, F. (2017). Draft Genome Sequence of the Polysaccharide-Degrading Marine Bacterium Pseudoalteromonas sp. Strain A601. Genome Announcements, 5(29). https://doi.org/10.1128/genomeA.00590-17

Li, X., Wang, Z., Lu, F., Zhang, H., Tian, J., He, L., … Tian, Y. (2018). Actinocorallia populi sp. nov., an endophytic actinomycete isolated from a root of Populus adenopoda (Maxim.). International Journal of Systematic and Evolutionary Microbiology, 68(7), 2325–2330. https://doi.org/10.1099/ijsem.0.002840

Lima, N. B., Gama, M. A. S., Mariano, R. L. R., Silva, W. J., Farias, A. R. G., Falcão, R. M., … Souza, E. B. (2017). Complete Genome Sequence of Xanthomonas campestris pv. viticola Strain CCRMXCV 80 from Brazil. Genome Announcements, 5(46). https://doi.org/10.1128/genomeA.01263-17

Lincoln, S. A., Hamilton, T. L., Valladares Juárez, A. G., Schedler, M., Macalady, J. L., Müller, R., & Freeman, K. H. (2015). Draft Genome Sequence of the Piezotolerant and Crude Oil-Degrading Bacterium Rhodococcus qingshengii Strain TUHH-12. Genome Announcements, 3(2). https://doi.org/10.1128/genomeA.00268-15

Loyola, D. E., Navarro, C., Uribe, P., García, K., Mella, C., Díaz, D., … Espejo, R. T. (2015). Genome diversification within a clonal population of pandemic Vibrio parahaemolyticus seems to depend on the life circumstances of each individual bacteria. BMC Genomics, 16(1), 176. https://doi.org/10.1186/s12864-015-1385-8

Maitra, N., Whitman, W. B., Ayyampalayam, S., Samanta, S., Sarkar, K., Bandopadhyay, C., … Manna, S. K. (2014). Draft Genome Sequence of the Aquatic Phosphorus-Solubilizing and -Mineralizing Bacterium Bacillus sp. Strain CPSM8. Genome Announcements, 2(1). https://doi.org/10.1128/genomeA.01265-13

Mashima, J., Kodama, Y., Kosuge, T., Fujisawa, T., Katayama, T., Nagasaki, H., … Takagi, T. (2016). DNA data bank of Japan (DDBJ) progress report. Nucleic Acids Research, 44(D1), D51–D57. https://doi.org/10.1093/nar/gkvll05

Mazuet, C., Legeay, C., Sautereau, J., Ma, L., Bouchier, C., Bouvet, P., & Popoff, M. R. (2016). Diversity of Group I and II Clostridium botulinum Strains from France Including Recently Identified Subtypes. Genome Biology and Evolution, 8(6), 1643–1660. https://doi.org/10.1093/gbe/evwl01

Mehrabadi, J. F., Mirzaie, A., Ahangar, N., Rahimi, A., & Rokni-Zadeh, H. (2016). Draft Genome Sequence of Kocuria rhizophila RF, a Radiation-Resistant Soil Isolate. Genome Announcements, 4(2). https://doi.org/10.1128/genomeA.00095-16

Naushad, S., Barkema, H. W., Luby, C., Condas, L. A. Z., Nobrega, D. B., Carson, D. A., & De Buck, J. (2016). Comprehensive Phylogenetic Analysis of Bovine Non-aureus Staphylococci Species Based on Whole-Genome Sequencing. Frontiers in Microbiology, 7. https://doi.org/10.3389/fmicb.2016.01990

Niehaus, E.-M., Kim, H.-K., Münsterkötter, M., Janevska, S., Arndt, B., Kalinina, S. A., … Tudzynski, B. (2017). Comparative genomics of geographically distant Fusarium fujikuroi isolates revealed two distinct pathotypes correlating with secondary metabolite profiles. PLOS Pathogens, 13(10), e1006670. https://doi.org/10.1371/journal.ppat.1006670

Norrander, J., Kempe, T., & Messing, J. (1983). Construction of improved M13 vectors using oligodeoxynucleotide-directed mutagenesis. Gene, 26(1), 101–106. https://doi.org/10.1016/0378-1119(83)90040-9

O’Leary, N. A., Wright, M. W., Brister, J. R., Ciufo, S., Haddad, D., McVeigh, R., … Pruitt, K. D. (2016). Reference sequence (RefSeq) database at NCBI: current status, taxonomic expansion, and functional annotation. Nucleic Acids Research, 44(D1), D733–D745. https://doi.org/10.1093/nar/gkv1189

Orsi, W. D., Edgcomb, V. P., Christman, G. D., & Biddle, J. F. (2013). Gene expression in the deep biosphere. Nature, 499(7457), 205–208. https://doi.org/10.1038/nature12230

Ortiz, E. M., Berretta, M. F., Navas, L. E., Benintende, G. B., Amadio, A. F., & Zandomeni, R. O. (2015). Draft Genome Sequence of Geobacillus sp. Isolate T6, a Thermophilic Bacterium Collected from a Thermal Spring in Argentina. Genome Announcements, 3(4). https://doi.org/10.1128/genomeA.00743-15

Parke, D. (1990). Construction of mobilizable vectors derived from plasmids RP4, pUC18 and pUC19. Gene, 93(1), 135–137. https://doi.org/10.1016/0378-1119(90)90147-J

Pavlov, M. S., Lira, F., Martinez, J. L., Olivares, J., & Marshall, S. H. (2015). Draft Genome Sequence of Antarctic Pseudomonas sp. Strain KG01 with Full Potential for Biotechnological Applications. Genome Announcements, 3(4). https://doi.org/10.1128/genomeA.00906-15

Peng, T., Pan, S., Christopher, L., Sparling, R., & Levin, D. B. (2015). Draft Genome Sequence of Thermoanaerobacter sp. Strain YS13, a Novel Thermophilic Bacterium. Genome Announcements, 3(3). https://doi.org/10.1128/genomeA.00584-15

Perron, A., Raymond, P., & Simard, R. (2006). The Occurrence of Antibiotic Resistance Genes in Taq Polymerases and a Decontamination Method Applied to the Detection of Genetically Modified Crops. Biotechnology Letters, 28(5), 321–325. https://doi.org/10.1007/s10529-005-5931-3

Pisarenko, S. V., Kovalev, D. A., Volynkina, A. S., Ponomarenko, D. G., Rusanova, D. V., Zharinova, N. V., … Kulichenko, A. N. (2018). Global evolution and phylogeography of Brucella melitensis strains. BMC Genomics, 19(1), 353. https://doi.org/10.1186/s12864-018-4762-2

Planet, P. J., Diaz, L., Kolokotronis, S.-O., Narechania, A., Reyes, J., Xing, G., … Arias, C. A. (2015). Parallel Epidemics of Community-Associated Methicillin-Resistant Staphylococcus aureus USA300 Infection in North and South America. Journal of Infectious Diseases, 212(12), 1874–1882. https://doi.org/10.1093/infdis/jiv320

Poehlein, A., Freese, H. M., Daniel, R., & Simeonova, D. D. (2014). Draft Genome Sequence of Serratia sp. Strain DD3, Isolated from the Guts of Daphnia magna. Genome Announcements, 2(5). https://doi.org/10.1128/genomeA.00903-14

Poehlein, A., Solano, J. D. M., Flitsch, S. K., Krabben, P., Winzer, K., Reid, S. J., … Dürre, P. (2017). Microbial solvent formation revisited by comparative genome analysis. Biotechnology for Biofuels, 10(1), 58. https://doi.org/10.1186/s13068-017-0742-z

Pore, S. D., Arora, P., & Dhakephalkar, P. K. (2014). Draft Genome Sequence of Geobacillus sp. Strain FW23, Isolated from a Formation Water Sample. Genome Announcements, 2(3). https://doi.org/10.1128/genomeA.00352-14

Pridmore, R. D. (1987). New and versatile cloning vectors with kanamycin-resistance marker. Gene, 56(2-3), 309–312. https://doi.org/10.1016/0378-1119(87)90149-1

Quigley, N. B., & Reeves, P. R. (1987). Chloramphenicol resistance cloning vector based on pUC9. Plasmid, 17(1), 54–57. https://doi.org/10.1016/0147-619X(87)90008-4

Register, K. B., Ivanov, Y. V., Jacobs, N., Meyer, J. A., Goodfield, L. L., Muse, S. J., … Losada, L. (2015). Draft Genome Sequences of 53 Genetically Distinct Isolates of Bordetella bronchiseptica Representing 11 Terrestrial and Aquatic Hosts: TABLE 1. Genome Announcements, 3(2). https://doi.org/10.1128/genomeA.00152-15

Reid, A. J., Blake, D. P., Ansari, H. R., Billington, K., Browne, H. P., Bryant, J., … Pain, A. (2014). Genomic analysis of the causative agents of coccidiosis in domestic chickens. Genome Research, 24(10), 1676–1685. https://doi.org/10.1101/gr.168955.113

Reuter, S., Connor, T. R., Barquist, L., Walker, D., Feltwell, T., Harris, S. R., … Thomson, N. R. (2014). Parallel independent evolution of pathogenicity within the genus Yersinia. Proceedings of the National Academy of Sciences, 111(18), 6768–6773. https://doi.org/10.1073/pnas.1317161111

Riojas, M. A., McGough, K. J., Rider-Riojas, C. J., Rastogi, N., & Hazbón, M. H. (2018). Phylogenomic analysis of the species of the Mycobacterium tuberculosis complex demonstrates that Mycobacterium africanum, Mycobacterium bovis, Mycobacterium caprae, Mycobacterium microti and Mycobacterium pinnipedii are later heterotypic synonyms of Mycob. International Journal of Systematic and Evolutionary Microbiology, 68(1), 324–332. https://doi.org/10.1099/ijsem.0.002507

Rodriguez, R. L. (1988). Vectors: a survey of molecular cloning vectors and their uses. Biotechnology (Reading, Mass.), 10, 1–578.

Shah, B., Jain, K., Patel, N., Pandit, R., Patel, A., Joshi, C. G., & Madamwar, D. (2015). Draft Genome Sequence of Paenibacillus sp. Strain DMB20, Isolated from Alang Ship-Breaking Yard, Which Harbors Genes for Xenobiotic Degradation. Genome Announcements, 3(3). https://doi.org/10.1128/genomeA.00554-15

Sharma, C., Kumar, N., Meis, J. F., Pandey, R., & Chowdhary, A. (2015). Draft Genome Sequence of a Fluconazole-Resistant Candida auris Strain from a Candidemia Patient in India. Genome Announcements, 3(4). https://doi.org/10.1128/genomeA.00722-15

Soggiu, A., Piras, C., Gaiarsa, S., Bendixen, E., Panitz, F., Bendixen, C., … Roncada, P. (2015). Draft Genome Sequence of Clostridium tyrobutyricum Strain DIVETGP, Isolated from Cow’s Milk for Grana Padano Production. Genome Announcements, 3(2). https://doi.org/10.1128/genomeA.00213-15

Song, J. S., Lee, J. H., Lee, J.-H., Jeong, B. C., Lee, W.-K., Lee, S. H. (2006). Removal of contaminating TEM-la beta-lactamase gene from commercial Taq DNA polymerase. Journal of Microbiology (Seoul, Korea), 44(1), 126–8. Retrieved from http://www.ncbi.nlm.nih.gov/pubmed/16554728

Tian, J., Xu, L., Zhang, S., Sun, W., Chu, X., & Wu, N. (2014). Draft Genome Sequence of the Organophosphorus-Degrading Bacterium Pseudomonas sp. Strain 1-7, Isolated from Organophosphorus-Polluted Sludge. Genome Announcements, 2(5). https://doi.org/10.1128/genomeA.00993-14

Vélez, J. R., Cameron, M., Rodriguez-Lecompte, J. C., Xia, F., Heider, L. C., Saab, M., … Sánchez, J. (2017). Whole-Genome Sequence Analysis of Antimicrobial Resistance Genes in Streptococcus uberis and Streptococcus dysgalactiae Isolates from Canadian Dairy Herds. Frontiers in Veterinary Science, 4. https://doi.org/10.3389/fvets.2017.00063

Versluis, D., D’Andrea, M. M., Ramiro Garcia, J., Leimena, M. M., Hugenholtz, F., Zhang, J., … Passel, M. W. J. van. (2015). Mining microbial metatranscriptomes for expression of antibiotic resistance genes under natural conditions. Scientific Reports, 5(1), 11981. https://doi.org/10.1038/srep11981

Vieira, J., & Messing, J. (1982). The pUC plasmids, an M13mp7-derived system for insertion mutagenesis and sequencing with synthetic universal primers. Gene, 19(3), 259–268. https://doi.org/10.1016/0378-1119(82)90015-4

Visnovsky, S. B., Fiers, M., Lu, A., Panda, P., Taylor, R., & Pitman, A. R. (2016). Draft Genome Sequences of 18 Strains of Pseudomonas Isolated from Kiwifruit Plants in New Zealand and Overseas: TABLE 1. Genome Announcements, 4(2). https://doi.org/10.1128/genomeA.00061-16

Wang, X., Tao, F., Gai, Z., Tang, H., & Xu, P. (2012). Genome Sequence of the Welan Gum-Producing Strain Sphingomonas sp. ATCC 31555. Journal of Bacteriology, 194(21), 5989–5990. https://doi.org/10.1128/JB.01486-12

Wattam, A. R., Davis, J. J., Assaf, R., Boisvert, S., Brettin, T., Bun, C., … Stevens, R. L. (2017). Improvements to PATRIC, the all-bacterial Bioinformatics Database and Analysis Resource Center. Nucleic Acids Research, 45(D1), D535–D542. https://doi.org/10.1093/nar/gkw1017

Yan, X., Ge, H., Huang, T., Hindra, Yang, D., Teng, Q., … Shen, B. (2016). Strain Prioritization and Genome Mining for Enediyne Natural Products. MBio, 7(6). https://doi.org/10.1128/mBio.02104-16

Youngblut, N. D., Wirth, J. S., Henriksen, J. R., Smith, M., Simon, H., Metcalf, W. W., & Whitaker, R. J. (2015). Genomic and phenotypic differentiation among Methanosarcina mazei populations from Columbia River sediment. The ISME Journal, 9(10), 2191–2205. https://doi.org/10.1038/ismej.2015.31

Zatuga, J., Stragier, P., Baeyen, S., Haegeman, A., Van Vaerenbergh, J., Maes, M., & De Vos, P. (2014). Comparative genome analysis of pathogenic and non-pathogenic Clavibacter strains reveals adaptations to their lifestyle. BMC Genomics, 15(1), 392. https://doi.org/10.1186/1471-2164-15-392

Zeil, C., Widmann, M., Fademrecht, S., Vogel, C., & Pleiss, J. (2016). Network Analysis of Sequence-Function Relationships and Exploration of Sequence Space of TEM β-Lactamases. Antimicrobial Agents and Chemotherapy, 60(5), 2709–2717. https://doi.org/10.1128/AAC.02930-15

Zeng, H., Zhang, J., Wu, Q., He, W., Wu, H., Ye, Y., … Xue, L. (2018). Reconstituting the History of Cronobacter Evolution Driven by Differentiated CRISPR Activity. Applied and Environmental Microbiology, 84(10). https://doi.org/10.1128/AEM.00267-18

Zhang, W., Lohman, A. W., Zhuravlova, Y., Lu, X., Wiens, M. D., Hoi, H., … Campbell, R. E. (2017). Optogenetic control with a photocleavable protein, PhoCI. Nature Methods, 14(4), 391–394. https://doi.org/10.1038/nmeth.4222

Zhurina, D., Dudnik, A., Waidmann, M. S., Grimm, V., Westermann, C., Breitinger, K. J., … Riedel, C. U. (2013). High-Quality Draft Genome Sequence of Bifidobacterium longum E18, Isolated from a Healthy Adult. Genome Announcements, 1(6). https://doi.org/10.1128/genomeA.01084-13

Zwick, M. E., Joseph, S. J., Didelot, X., Chen, P. E., Bishop-Lilly, K. A., Stewart, A. C., … Read, T. D. (2012). Genomic characterization of the Bacillus cereus sensu lato species: Backdrop to the evolution of Bacillus anthracis. Genome Research, 22(8), 1512–1524. https://doi.org/10.1101/gr.134437.111

